# Coppice conversion of European beech (*Fagus sylvatica* L.): natural evolution, periodic thinning, regeneration cutting

**DOI:** 10.1101/2025.11.24.690100

**Authors:** Francesco Chianucci, Giada Bertini, Gioele Tiberi, Giada Lazzerini, Luca Marchino, Maurizio Piovosi, Andrea Cutini

**Author notes:** Corresponding author (Francesco Chianucci), (Giada Bertini).

## Abstract

Beech coppice forests have shaped the mountainous European landscape for centuries. The socio-economical changes occurred over the last 60 years have led to a progressive decline in coppice management, which has resulted in either the abandonment of traditional coppice management, or into active coppice conversion to high forests. Given the long-term process of this process, an ecological, long-term perspective is required to understand the ecological implication of different management practices in these forests.

We investigated the influence of conversion management on canopy attributes (leaf litter and seed production, leaf area index). The management options considered were traditional conversion management, based on periodic thinning, and alternative conversion, based on anticipated seed cutting and final harvesting, which were compared against natural evolution (unthinned control). Results showed that the differences between natural evolution and traditional conversion were largest in the years immediately after thinning, and then reduced with full canopy (leaf litter and leaf area index) recovery after 10 years. Conversely, the alternative method with the anticipated seed cutting significantly enhanced canopy heterogeneity and further accelerates the transition to high forest, with dense beech saplings reaching an height over 8 meters eight years post-harvest.

## 1. Introduction

Coppice forests have shaped the European landscape for centuries. Currently, coppice forests cover about 23 million hectares in the Mediterranean area (McGrath et al. 2015), still representing a significant part of European forests (Forest Europe 2020).

In Mediterranean areas, beech (*Fagus sylvatica* L.) forests have a long history of coppicing, which traditionally sustained the economy of people living in mountainous areas, providing a sustainable source of fuelwood, timber, and other forest products up to the mid-20th century (Fabbio 2016). Such intensive use has significantly modified the distribution, composition and structure of these forests (Fabbio 2016; Antonucci et al. 2021). The socio-economical changes occurred over the last 60 years have led to a progressive decline in coppice management in this species (Nocentini 2009). Such trend has resulted in either the abandonment of traditional coppice management, where former coppice stands were left to develop naturally, or into active management conversion towards high forests (Cutini et al. 2015). Anyway, a low diversification both at stand and landscape level over large surfaces occurred, which resulted into the need to define sustainable management and planning option to increase structural heterogeneity in these forests (Cutini et al. 2021).

Active conversion to high forest is typically carried out by progressively reducing tree density with periodic thinning, with the first intervention carried out soon at the end of the coppice rotation (Amorini et al 1990; Amorini and Fabbio 1986, 1990). Conversion is then achieved with seedling establishment after the last regeneration cutting, according to the uniform shelterwood system. The rotation period of this conversion system is 100– 150 years, depending on soil fertility, but the regeneration stage has rarely been filled in practice (Cutini et al. 2015). Accordingly, most former beech coppice stands are currently in a transition stage towards high forest (Nocentini 2009; Cutini et al. 2009; Mattioli et al. 2015). Whatever the management applied, such transition is a long-term, and thus poorly documented, process Understanding the influence of management on these forests is also relevant for designing adaptive management strategies to cope with climate changes, which is a priority in modern climate-smart forestry, particularly for these forests, which are relatively less studied (Jump, Hunt, and Peñuelas 2006).

In this contribution, we reported results of a long-term silvicultural trial aimed at evaluating the influence of coppice conversion in mountainous beech forests in Northern Apennines, Italy. The trial started in 1972 in beech coppice stands at the end of the rotation period, and involved two active conversion schemes:

- The traditional conversion based on periodic medium-heavy thinnings, carried out every 15-20 years to progressively reduce the density towards the future high forest composition (Chianucci, Salvati, et al. 2016, Bertini et al., submitted)
- An alternative conversion based on periodic thinning, followed by an anticipated seed cutting in stands aged 65 years (Cutini et al. 2015), and the final regeneration cutting 14 years afterwards (this study).

Both active conversion schemes were compared with a stand left to develop naturally (unthinned control). Previous studies indicated that active conversion accelerate the transition towards more mature, high forest structure, compared to natural evolution pattern in these forests, after 40 years from beginning the conversion (Chianucci, Salvati, et al. 2016) and ten years after the anticipated seed cutting (Cutini et al. 2015). In the current study, we extended the previous findings to address these specific questions:

- What is the influence of active management 50 years after the beginning of conversion?
- How the conversion is achieved after final cutting, compared to the traditional conversion and natural evolution patterns?

We used an ecological approach, involving evaluating the influence of management on annual canopy structure (leaf area index, litter production and its partitioning), which were measured every year. To the best of our knowledge, this is the first study reporting a complete coppice conversion in Mediterranean beech forests.

## 2. Material and methods

### 2.1. Study area and experimental trial

The study was carried out in Alpe di Catenaia, a moutainous forest in Northern Apennines (43° 49N; 11° 49 E; Figure 1).

**Figure 1.**
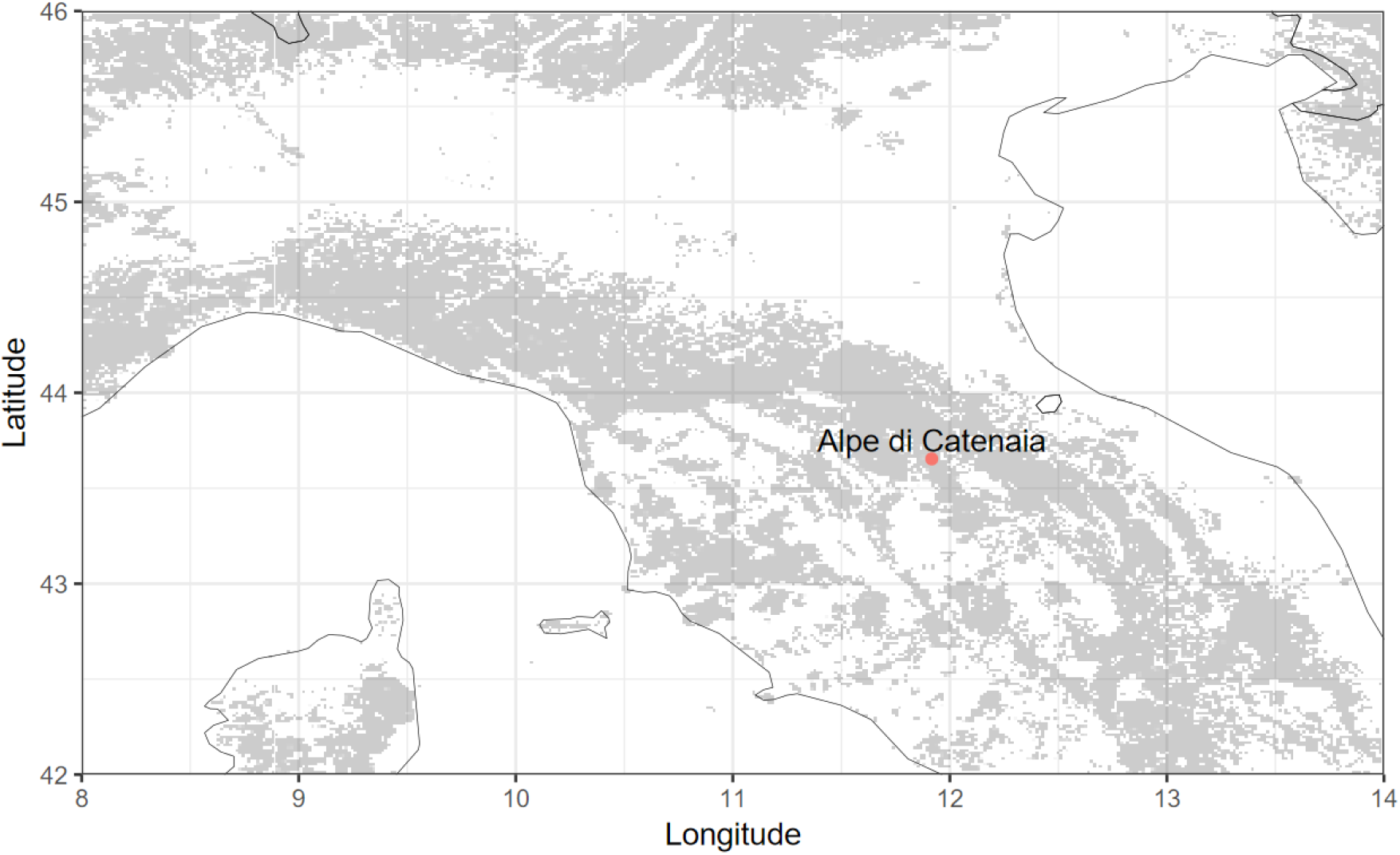
Study area. Grey areas represents forests with a tree cover greater than 40% according to the map of Kempeneers et al. 2011.

The climate is temperate, with dry summer and rainy winters. The altitude ranged between 400 and 1400 m, where beech forests dominated at altitudes above 900 m. The experimental trial included three permanent plots each located in 1 ha stands established in 1972 (Amorini et al. 2010), representing the natural evolution pattern and two active conversion systems.

The active conversion scheme consists in gradually reducing the number of shoots by periodic, medium-heavy thinnings from below (Ciancio et al. 2006; Amorini et al. 2010; Bertini et al., submitted). The interval between consecutive thinning allows a temporary opening of the canopy cover, to enable seed regeneration under increasing below-canopy light availability. In the uniform shelterwood system, regeneration cuttings are the final step of this conversion scheme, which consists of a first seed cutting, which is a strong thinning with retention of large, seed-bearer trees, an optional second thinning, and the final regeneration cutting. The regeneration period between the first seed cutting and the last regeneration cutting is typically 10-30 years, to allows seed production and seedling establishment in the intervening period.

### 2.2 Silvicultural treatment

The three plots were selected in stands managed as coppice with standards, aged 27 years in 1972, i.e. at the end of the rotation period. In 1972, a first thinning was undertaken in two stands to actively prepare them to conversion to high forest. Another stand was left to develop naturally since 1972.

In the traditional conversion system, four thinnings, of medium-heavy intensity, were carried out in 1972, 1987, 2002 and 2019, which progressively reduced standing basal area by 29%-44%. Compared with natural evolution, this traditional conversion system is profitable due to combination of frequent harvest and the development of more growth-efficient trees (Cutini et al. 2015; Bertini et al. submitted). Periodic thinning contributes to accelerate the transition of these managed stands towards more adult stand conditions, as larger growing space allows larger crown enlargement of retained trees, supporting seed production and below-canopy light to favor seed regeneration (Cutini et al. 2015; Chianucci, Salvati, et al. 2016). Detailed stand structure data are available in Amorini et al. 2010 and Bertini et al. (submitted).

In the alternative active conversion scheme, two low thinnings of medium-heavy intensity were carried out in 1972 and 1987, which were comparable with those undertaken with the traditional conversion type. In September 2002, at the age of 65, an anticipated seed cutting was carried out reducing the stand basal area by 53% (Cutini at al. 2010). The seed cutting produced stronger modification in canopy structure, improving growth efficiency because of higher resource availability, supporting higher seed production, which further accelerated the progression of the stand towards more adult stand conditions, compared with traditional management (Cutini et al. 2015). After ten years from anticipated seed cutting, vegetation sampling showed that beech saplings on average reached 1.5 m, and understory cover was about 40%, while in traditional thinning the average sapling height was 0.4 m, with an understory cover of 15% (Cutini et al. 2015). As the beech regeneration after 14 years from seed cutting were above the deer browsing height ( 1.8 m (Chianucci, Mattioli, et al. 2015)), and the tree crown almost fully recovered (Chianucci, Disperati, et al. 2016), final harvesting was carried out in 2016, to remove all the overstory trees to release the established natural beech regeneration.

### 2.3 Litterfall measurements

Litter production and its partitioning were estimated using littertraps, as proposed in (Chianucci and Cutini 2013) and following described. Based on stand homogeneity, nine to fifteen traps, each 0.25 in size, were systematically placed in each stand, with trap spaced 7-20 m. Litter was collected from 1992 to 2024 (2003-2016 in the alternative conversion scheme, after seed cutting and before regeneration cutting) every year in fall-winter every two weeks, with the last collection soon after the last litter fall. Litter was then separated in laboratory into leaf, woody and reproductive (husks and seeds) components, and then dried 24 h in a fan-forced oven, at 85° 2° C. The annual dry mass (Mg ha^-1^) of litter components was then calculated for each stand.

### 2.4 Leaf area index measurements

Leaf area index (LAI) was estimated using the Licor LAI 2000/2200 Plant Canopy Analyzer (Licor, NE, US), following the method described in Chianucci, Macfarlane, et al. (2015). Below-canopy measurements were collected in summer, in diffuse sky conditions, from 1999 to 2025 (2002-2016 in the alternative conversion scheme, before the seed and regeneration cuttings), along the same grid used for litter traps. An above-canopy reference was collected in a clearing close to each stand. The sensor was covered by a 90° view-cap to avoid influence of operator and surrounding trees on measurements. We considered the effective leaf area index (*Le*), which was calculated using a linear averaging gap fraction approach (Chianucci 2019). An apparent clumping factor (ACF) was also calculated and used as a proxy of canopy heterogeneity. ACF calculate the deviation from a random (Poisson) distribution of foliage, and varied between 0 (maximum heterogeneity) to 1 (maximum homogeneity).

### 2.5 Beech sapling regeneration

In 2024, a field sampling was carried out in the alternative conversion stand after 8 years from the final harvesting. Three circular sub-plots, each 5 m in radius were randomly placed in the stands, and all tree saplings (diameter ) measured. Tree height was also measured for 25% of saplings.

### 2.6 Statistical analyses

We compared the influence of different management regimes on canopy structure using one-Kruskall-Wallis. The trend of canopy attributes after thinning was also evaluate using Spearman’s *r* correlation.

## 3. Results

Figure 2 reports results from the litterfall sampling. In the traditional conversion scheme, thinning reduced leaf litter by 40% in 2002, from 4.56 to 2.75, and by 30% in 2019, from 2.77 to 1.96 .

**Figure 2.**
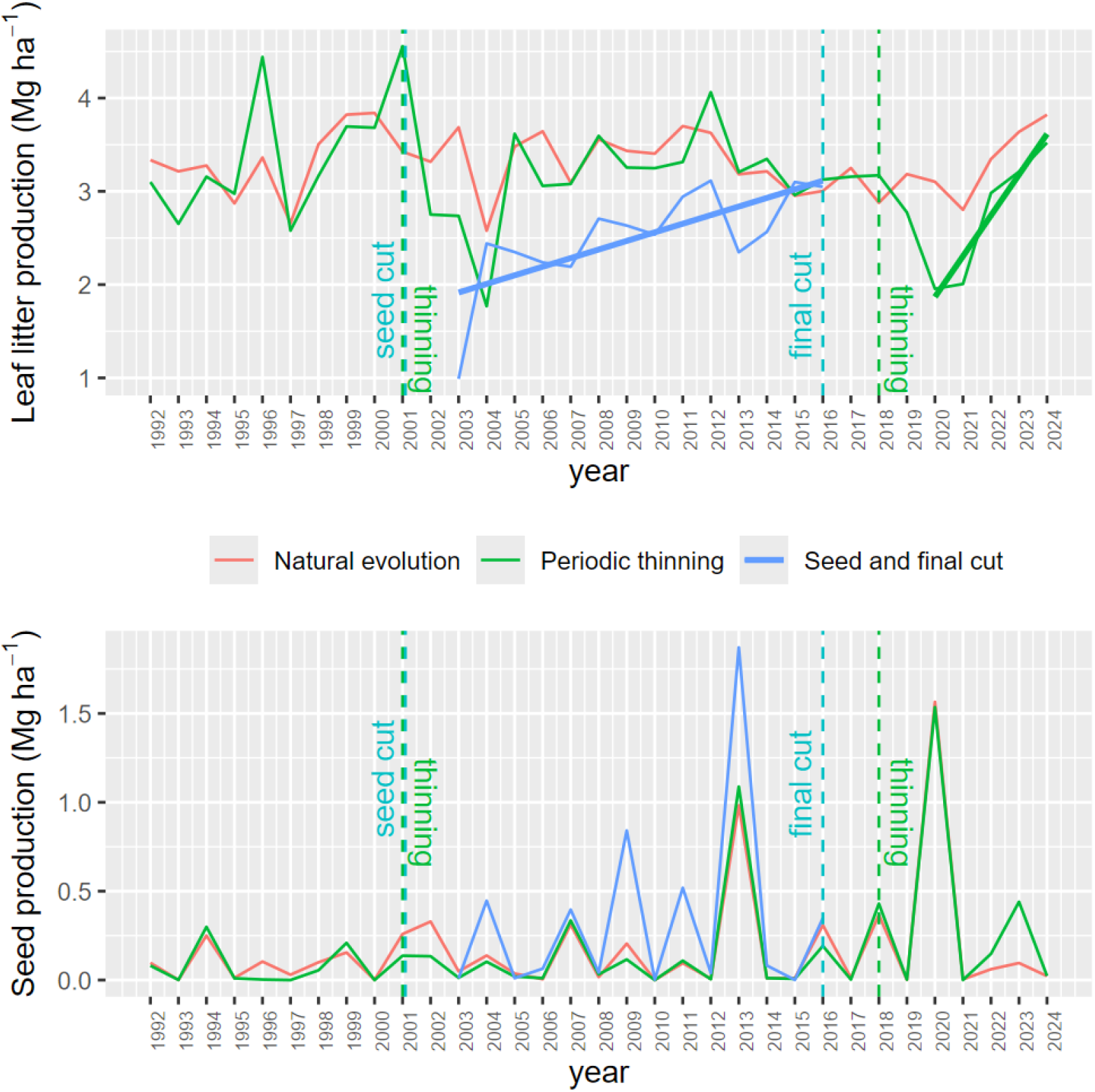
Trend of leaf and seed production in the three management options. Top: leaf litter production. Bottom: seed production. The lines indicate significant trend over time.

The trend of leaf production is not significant in the natural evolution stand, while it is significant in the alternative conversion, and after the second thinning in the traditional conversion (Spearman’s *r, p* < 0.05).

When considering seed production, the annual pattern is similar between the stands, and the differences are not significant (Kruskall-Wallis test, *p* > 0.05), although the active management trials, and particularly the alternative conversion, have larger seed production than natural evolution stand. Average (± standard error) seed production was 0.334 ± 1.14 in alternative conversion, 0.176 ± 0.06 in traditional conversion and 0.169 ± 0.05 in natural evolution stand. There is no significant trend in seed production over time (Figure 2).

In the alternative conversion scheme, anticipated seed cutting reduced leaf area index by 81% in 2002, from 5.2 to 1.0 (Figure 3). The trend of *Le* was not significant in the natural evolution stand, while it showed a significant increase in the alternative conversion, and in the period after the first and before the second thinning in the traditional conversion scheme (Spearman’s *r, p* < 0.05).

**Figure 3.**
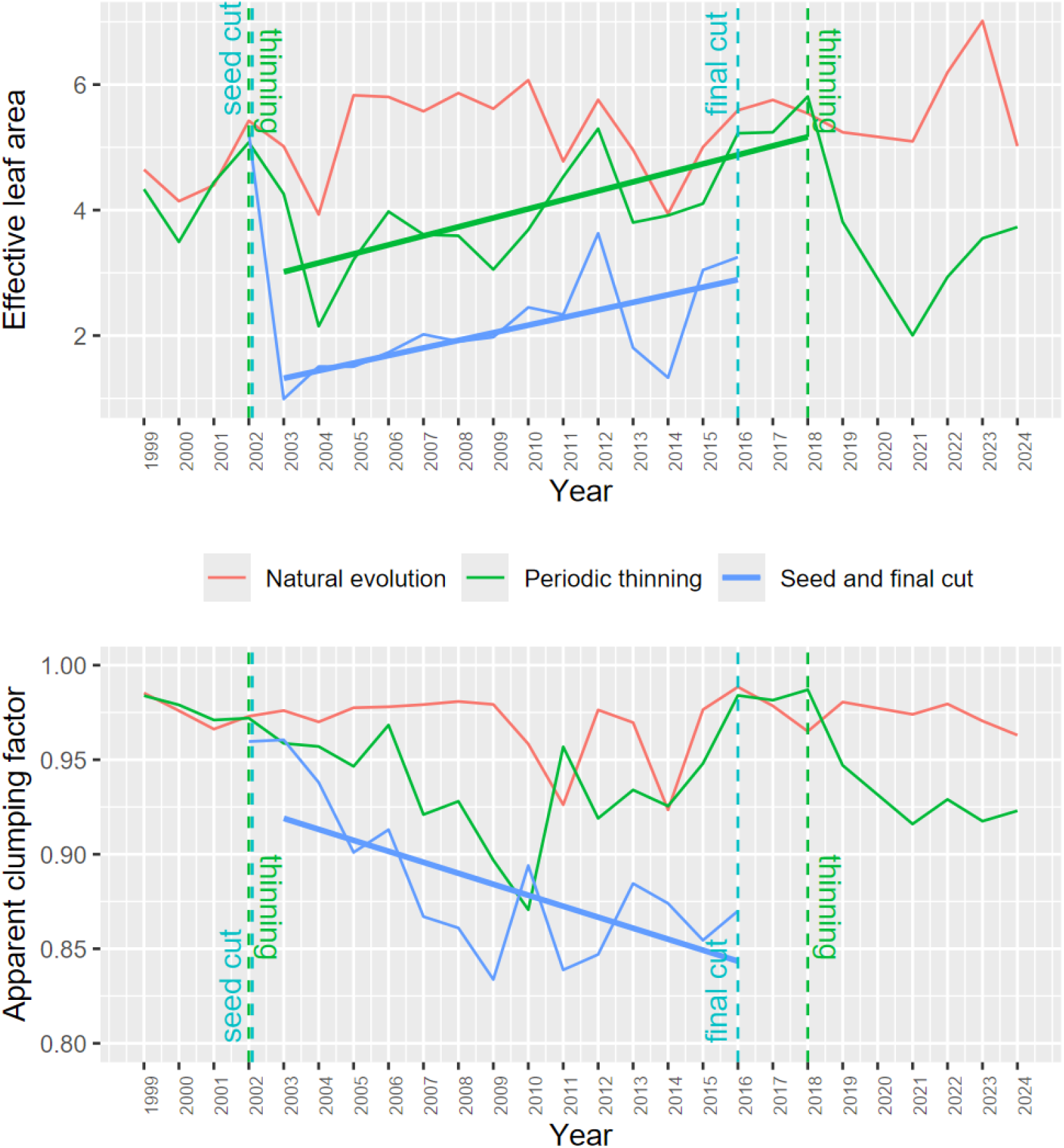
Trend of leaf area index and apparent clumping factor in the three management options. Top: Effective leaf area index (Le; m^2^ m^-2^). Bottom: Apparent clumping factor (ACF). The lines indicate significant trend over time.

The seed cutting also significantly increased the hetereogenity in the alternative conversion scheme, as proxied by ACF, which also showed a significant decrease over time (Spearman’s *r, p* < 0.05), while the trend is not significant for both natural evolution and traditional conversion schemes (Figure 3).

Finally, in the regeneration plots we found 434 beech saplings and 11 hop hornbeam (*Ostrya carpinifolia*). We found that after 8 years from final harvesting, the beech sapling is very dense and established, with a dominant height above 8 m (Table 1). We found that after 8 years from final harvesting, the beech sapling is very dense and established, with a dominant height above 8 m (Table 1).

**Table 1.** Mean beech sapling attributes in the regeneration plots. D: diameter at breast height (cm); H= height (m); G: plot basal area (m^2^ ha^-1^). SR: plot sapling density (n ha^-1^)

| attribute | mean | sd |
| --- | --- | --- |
| D | 3.19 | 0.68 |
| H | 8.22 | 0.67 |
| G | 14.16 | 3.08 |
| SR | 18420 | 4630 |

## 4. Discussion

This is the first study documenting a complete beech coppice conversion process, from first thinning to the last regeneration cutting. Our results agreed with previous findings, which indicated that anticipating seed and final harvesting can accelerate coppice transition to high forest in beech coppice stands aged 60-70 years (Cutini et al. 2010; Cutini et al. 2015). We further extend the previous results by indicating that after 8 years from final harvesting, the stand has a very dense and established beech regeneration, which confirmed the successful transition to high forests.

Conversely, the differences between traditional conversion and natural evolution stands were largest in the period after thinning, due to the canopy recovery (leaf litter and leaf area index), but reached a plateau between two thinning, leading to a slower progression of these stands towards high forests, as compared with the alternative conversion scheme (Cutini et al. 2015). In this line, a ten-year period is appropriate for fully canopy recovery after medium-heavy thinning, which makes traditional conversion scheme profitable due to frequent harvesting and intermediate volumes (Chianucci, Salvati, et al. 2016; Bertini et al., submitted).

A key strength of the proposed study is that we used a long-term perspective, involving use of canopy structure data that allows an ecological evaluation on the different coppice conversion management options. The obtained LAI and leaf litter values were in accordance with other mature beech stands (Değermenci et al. 2025; Meier and Leuschner 2008; Leuschner et al. 2006), which confirmed the sampled stands have medium-high productivity, which is pivotal for achieving successful conversion.

Interestingly, we did not find a positive trend in seed production in these stands over time, despite seed production should increase with age (Genet, Breda, and Dufrene 2009). However, the seed production values observed in two masting events (2013 and 2020) indicated very high production, which is comparable with hard mast in high forests beech stands in Europe (Chianucci et al. 2021; E Silva et al. 2012; Overgaard, Gemmel, and Karlsson 2007), confirming that medium-high productive stands can support steady seed production, which is a requisite for coppice conversion. We also found very frequent alternation of full-mast (e.g., 2013, 2020) and half-mast (e.g. 2009, 2016) events, which may confirm an observed continental trend of masting breakdown in this species, due to lower synchrony and fewer failure years, due to climate change (Bogdziewicz et al. 2020). Additionally, despite the differences in seed production were not significant in the stands, the stand after seed cutting showed higher seed production in mast years, compared with the other two management options. In this line, the three medium-hard masting events occurred in 2009, 2012 and particularly in 2013, may have likely further supported beech regeneration before final harvesting in this stand. As the climatic changes can reduce the reproductive potential of this species (i.e. increasing number of empty seeds; Bogdziewicz et al. 2023), this may suggest that the timing for seed cutting should be synchronized with hard mast events, particularly in situation where an already established beech regeneration is observed in actively managed stands.

Considering that most former beech coppice stands in Apennines are currently in a transition stage, the choice of adopting different management options at landscape scale can allows to increase landscape diversity, supporting the structural diversification of these previously uniformly managed stands, favoring faster transition to more mature stand structure, while achieving the multiple needs and goods that are demanded by these forests (Cutini et al. 2021; Chianucci et al. 2024).

## Acknowledgements

FC was supported by the National Recovery and Resilience Plan (NRRP), Mission 4 Component 2 Investment 1.4 - Call for tender No. 3138 of 16 December 2021, rectified by Decree n.3175 of 18 December 2021 of Italian Ministry of University and Research funded by the European Union - NextGenerationEU; Award Number: Project code CN_00000033, Concession Decree No. 1034 of 17 June 2022 adopted by the Italian Ministry of University and Research, CUP B83D21014060006 and CUP: J33C22001190001, Project title “National Biodiversity Future Center - NBFC”.

